# Validity of measures of reactive balance training intensity

**DOI:** 10.1101/2025.08.06.668970

**Authors:** Avril Mansfield, Sarah Thompson, Dario Luis Felipe Puentes Sanchez, David Jagroop, George Mochizuki

**Affiliations:** KITE-Toronto Rehabilitation Institute, University Health Network, Toronto, ON, Canada; Department of Physical Therapy, University of Toronto, Toronto, ON, Canada; Evaluative Clinical Sciences, Hurvitz Brain Sciences Program, Sunnybrook Research Institute, Toronto, ON, Canada; Department of Kinesiology, University of Waterloo, Waterloo, ON, Canada; Physiotherapy, National University of Colombia, Bogotá, Colombia; School of Kinesiology and Health Science, York University, Toronto, ON, Canada

**Keywords:** Postural balance, Kinematics, Arousal, Perception, Exercise

## Abstract

Intensity is a key component of exercise prescription. However, balance training intensity is not well defined. Consequently, balance training studies often do not report exercise intensity, which hinders implementing these interventions and developing exercise guidelines. This study aimed to validate measures of reactive balance training intensity. Healthy young (n=16) and older (n=15) adults experienced moving platform balance perturbations of varying magnitudes. Candidate intensity measures were: number of reactive steps; centre of mass displacement, velocity, and acceleration, and margin of stability 100 ms post-perturbation; peak electrodermal response; and OMNI Perceived Exertion Scale and Balance Intensity Scale scores. Correlations were examined between perturbation magnitude and candidate intensity measures. We also compared candidate intensity measures between backward- and forward-fall perturbations and between age groups using generalized linear mixed models. Peak centre of mass acceleration 100 ms after the perturbation was significant positively correlated with perturbation magnitude, with no significant age-group or direction effects, suggesting that this is a valid measure of absolute perturbation intensity. Number of reactive steps, peak electrodermal response, and OMNI Perceived Exertion Scale were significantly positively correlated with perturbation magnitude, and were significantly higher for backward-fall compared with forward fall perturbations and for older adults compared to younger adults at the same magnitude, suggesting that these are valid measures of relative perturbation intensity. These measures could be used to prescribe and report intensity of reactive balance training in future studies.

## INTRODUCTION

Frequency, Intensity, Time, and Type (i.e., FITT) is a commonly-used framework for prescribing exercise (1). Intensity is the challenge of exercise, which can be absolute or relative to a person’s maximal capacity (2). Relative exercise intensity is used for individualized exercise prescription so that intensity is within the person’s capabilities but challenging enough to stimulate adaptation (1). Unlike other forms of exercise, balance training lacks a clear approach to define or measure intensity (3–5). As a result, intensity is rarely or erroneously reported in balance training studies (2,4,6), hindering clinical implementation (5,7,8).

Reactive balance training (RBT) involves having participants experience repeated balance perturbations to evoke and practice balance reactions, such as reactive stepping (6,9). Balance perturbations are often delivered in studies using devices such as a moving platform (6), where perturbation magnitude, or absolute intensity, can be well controlled. For example, absolute intensity is directly defined by peak platform acceleration (10,11), which is proportional to the force of the balance perturbation. However, this approach to specifying intensity cannot be applied to RBT approaches that do not use moving platforms, and cannot be directly used to prescribe relative intensity. Therefore, a standardized intensity definition that can be applied to various RBT approaches is needed.

Proxy intensity measures are often used for other types of exercise, as it is not always feasible to measure exercise intensity directly (2). Candidate proxy intensity measures for RBT include behavioural response (e.g., reactive stepping), centre of mass (COM) motion, electrodermal activity, and participant subjective ratings. Reactive stepping is more prevalent for high compared to low-magnitude perturbations, and among balance-impaired compared to non-balance-impaired populations (12–14). Therefore, characterizing stepping reactions to perturbations could be one way to set relative RBT intensity (15–18), but can only distinguish broad ranges of intensity (e.g., ‘low’, ‘moderate’, or ‘high’). Peak centre of mass (COM) displacement and velocity 100 ms after perturbation onset, prior to onset of the balance reaction, was previously used to determine if two different balance perturbation types (waist pull and moving platform) were of similar magnitude (19). COM displacement and velocity were lower for waist pull compared to surface translation perturbations, but not different between older and young adults (19). As fewer multi-step reactions were observed for waist pulls compared to surface translations (19), these findings suggest that COM motion reflects absolute perturbation intensity. Margin of stability (MOS) is the distance between the extrapolated COM (calculated using COM displacement and velocity) and the edge of the base of support (20). The MOS indicates how close a person is to falling (21). Minimum MOS predicts stepping reactions after a perturbation (13), so may reflect relative RBT intensity. Balance perturbations evoke electrodermal responses (EDRs), which are phasic increases in electrodermal activity that begin ∼2 s and peak 4-5 s post-perturbation (22). EDR peaks were higher when responses to balance perturbations were restricted, and were therefore more challenging (23), and higher for people with stroke compared to healthy controls (24). These findings suggest that electrodermal activity reflects relative RBT intensity. The Balance Intensity Scale (BIS) is an intensity rating scale specific to balance tasks (25,26) that correlates with challenge of anticipatory balance training (25). The OMNI Perceived Exertion Scale (PES) is a more general rating of perceived exertion during exercise (27); OMNI PES scores correlate with challenge of anticipatory balance training (28).

We aimed to establish validity of measures of absolute and relative RBT intensity among healthy young and older adults. We tested eight candidate intensity measures: number of steps; peak COM displacement, velocity, and acceleration, and minimum MOS 100 ms post-perturbation; EDR; and BIS, and OMNI PES scores. Specific objectives were: 1) to evaluate correlations between candidate intensity measures and perturbation magnitude; and 2) to determine if candidate intensity measures differ between forward- and backward-directed perturbations and between groups of young and older adults at the same magnitude. Absolute RBT intensity measures will be strongly related to perturbation magnitude, with no evidence of direction or group effects. As the COM is closer to the back of the foot during standing, perturbations that cause backward motion are more challenging than those that cause forward motion, and older adults are expected to have worse reactive balance control than young adults. Therefore, perturbations at the same magnitude will be experienced as at higher intensity for backward than forward perturbations, and for older adults compared to young adults, and measures that differ between directions and groups will represent relative RBT intensity.

## METHODS

### Participants

Participants were healthy community-dwelling young (20-35 years old) and older adults (60-75 years old). Participants were excluded if they: were unable to stand independently for ≥1 min; were unable to walk independently for ≥10 m; had neurological conditions; had osteoporosis, lower extremity joint replacement or joint fusion, or recurrent dizziness; and/or had cognitive impairment (Montreal Cognitive Assessment (29) score <26). Participants were recruited between 20 July 2023 and 1 May 2026. Participants provided written informed consent prior to starting data collection. The study was approved by the University Health Network Research Ethics Board (study ID: 23-5382).

### Apparatus

Data collection occurred in a lab with a custom-built 6m x 3m motion platform. The platform can be precisely controlled to translate in the transverse plane with maximum acceleration, velocity, and displacement of 10 m/s^2^, 2 m/s and 2 m, respectively. The platform is equipped with a robotic safety harness gantry that allows participants to move freely, but prevents them from falling if they were unable to regain balance. An accelerometer (Series 7523A, Dynamic Transducer and Systems, Chatsworth, California, USA) placed on the platform recorded platform acceleration. Electrodermal activity was recorded wirelessly via disposable silver/silver chloride electrodes placed on the fingers (BioNomadix module, BN-PPGED-T, BIOPAC Systems, Inc., Goleta, California, USA). Nine to 12 motion capture cameras (Vicon Mx 40+ and MX F20 or Vicon Vero 2.2, Vicon Motion Capture Systems Ltd., Oxford, UK) were positioned around the room to capture kinematic data.

Motion capture data were sampled at 100 Hz. To collect electrodermal responses (EDR), the palmar surfaces of the index and middle fingers were cleaned with alcohol wipes. EDR and accelerometer data were sampled at 1,000 Hz. Data were stored for offline processing.

### Procedure

At the start of the session, the Montréal Cognitive Assessment (29) was completed to confirm eligibility. To describe general balance abilities of the study cohort, participants completed the mini-Balance Evaluation Systems Test (30), and Activities-specific Balance Confidence scale (31).

Participants were asked to wear comfortable tight-fitting athletic clothing and low-heel running shoes. Participants were outfitted with 67 reflective markers (46 single markers and 21 rigid-plate marker clusters). They stood in a standardized foot position (32) on the platform. Behavioural and physiologic responses differ between initial and later perturbations (33,34); therefore, participants first completed a familiarization block of trials consisting of 5 large-magnitude perturbations (peak acceleration: 3 m/s^2^) to mute these first-trial effects. Participants then experienced 80 (older adults) or 144 (young adults) multi-directional perturbations (forward, backward, left, right) of varying intensities (Table 1); a shorter protocol was used with older adult participants to avoid fatigue. A range of intensities that should evoke no-step to multi-step reactions, but not falls, was chosen (9,14). Perturbations were delivered in four blocks, with one perturbation per direction per intensity level in each block. There was a 20-30 s break between consecutive perturbations to allow electrodermal activity to return to baseline (22). Rest breaks were scheduled between blocks, and participants could request rest breaks at any time. Perturbations were presented in an unpredictable sequence to prevent participants from planning responses. Additionally, to prevent participants from learning to use the deceleration phase of the perturbation waveform to regain stability (35), two different waveform shapes were used: one with the deceleration phase equal and opposite to the acceleration phase, and one with a deceleration phase twice as long (and, therefore, with a reduced peak) as the acceleration phase (13). Participants were instructed to react naturally, but, if they stepped, to try to take as few steps as possible.

**Table 1:** Perturbation characteristics. The acceleration pulse was delivered over 300ms.

| Intensity level | Peak acceleration (m/s<sup>2</sup>) | Peak velocity (m/s) | Total displacement (m) | Group |
| --- | --- | --- | --- | --- |
| 1 | 1 | 0.3 | 0.090 | Both |
| 2 | 1.25 | 0.375 | 0.113 | YA only |
| 3 | 1.5 | 0.45 | 0.135 | Both |
| 4 | 1.75 | 0.525 | 0.158 | YA only |
| 5 | 2 | 0.6 | 0.180 | Both |
| 6 | 2.25 | 0.675 | 0.203 | YA only |
| 7 | 2.5 | 0.75 | 0.225 | Both |
| 8 | 2.75 | 0.825 | 0.248 | YA only |
| 9 | 3 | 0.9 | 0.270 | Both |

YA = young adults.

Participants were asked to score the effort of responding to the perturbation on the 5-point Balance Intensity Scale (BIS; 25) for one half of the perturbations, and using the 11-point OMNI PES (27,28) for the other half of the perturbations. The order of presentation of the BIS and OMNI Perceived Exertion Scale was counter-balanced across participants.

### Data processing

Kinematic data were filtered using a zero-phase lag 2^nd^ order low-pass Butterworth filter at 6 Hz. Whole-body COM position was calculated from kinematic data as the weighted average of individual body segments, using sex- and age-specific COM models (36,37). COM velocity and acceleration were calculated from COM position. MOS was calculated relative to the appropriate base of support boundary (e.g., toe for perturbations that cause forward COM motion;,38) using the following formulae (20):

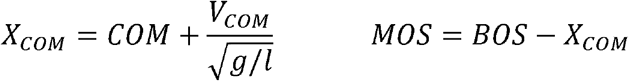

where *X*_*COM*_ is extrapolated COM, *COM* is COM position, *V*_*COM*_ is COM velocity, *g* is gravitational acceleration, *l* is the pendulum length, *MOS* is margin of stability, and *BOS* is the base of support boundary position. The MOS sign was adjusted, as appropriate, to yield a negative value when the X_COM_ was outside the base of support (38). Perturbation onset time was defined as the time when the platform acceleration exceeded 0.1 m/s^2^. Peak COM position, velocity, and acceleration, and minimum MOS 100 ms post-perturbation were calculated. Electrodermal data were filtered using a zero phase lag 2^nd^ order low-pass Butterworth filter at 2 Hz. Baseline electrodermal level was the mean electrodermal activity 1 s prior to the perturbation. EDR was the difference between maximum electrodermal value in the 10 s post-perturbation and the baseline electrodermal level (23).

### Data analysis

Poisson regression (number of steps) or linear regression (all other candidate measure) with generalized estimating equations and repeated measures (to account for multiple observations per participant;,39) were used to determine the relationship between perturbation magnitude and each candidate measure (number of steps; EDR amplitude; minimum MOS; peak COM displacement, velocity, and acceleration; and BIS and OMNI Perceived Exertion Scale scores) for each age group separately.

As the COM tends to be closer to the back than the front of the foot (40,41) perturbations that induce backward COM motion are more challenging than perturbations that induce forward motion. Furthermore, as older adults tend to have worse reactive balance control than young adults, older adults should experience perturbations at the same magnitude as a higher relative intensity than young adults. We therefore examined group and perturbation direction differences to test whether the candidate intensity measures reflect relative intensity, versus perturbation magnitude alone (absolute intensity), using generalized linear mixed models, with fixed effects of group, direction, and the interaction of group with direction, a random effect of participant, and magnitude included as a confounding variable. When there were significant effects, effect sizes for pairwise comparisons were presented mean difference with 95% confidence intervals in square brackets. Alpha was 0.05 for all analyses.

## RESULTS

Thirty-two participants (16 young adults and 16 older adults) were recruited. One older adult participant withdrew from the study after the familiarization block. Therefore, 15 older adult participants were included in the analysis. Participant characteristics are included in Table 2.

**Table 2:** Participant characteristics. Values presented are means with standard deviations in parentheses for continuous variables, or counts for categorical variables.

|  | Young adults | Older adults |
| --- | --- | --- |
| Age (years) | 24.9 (3.3) | 67.9 (3.6) |
| Sex |  |  |
| Female (number) | 8 | 7 |
| Male (number) | 8 | 8 |
| Height (m) | 1.74 (0.10) | 1.70 (0.09) |
| Mass (kg) | 72.8 (17.0) | 65.5 (11.8) |
| Mini-BEST score | 26.7 (1.1) | 26.4 (1.2) |
| ABC score (%) | 97.6 (2.0) | 97.0 (3.2) |

ABC = Activities Specific Balance confidence; mini-BEST = mini-Balance Evaluation Systems Test

There were significant negative (MOS) and positive (all other measures) relationships between perturbation magnitude and all measures (p<0.0001 for all measures, except EDR amplitude for young adults: p=0.0004; Figure 1).

**Figure 1:**
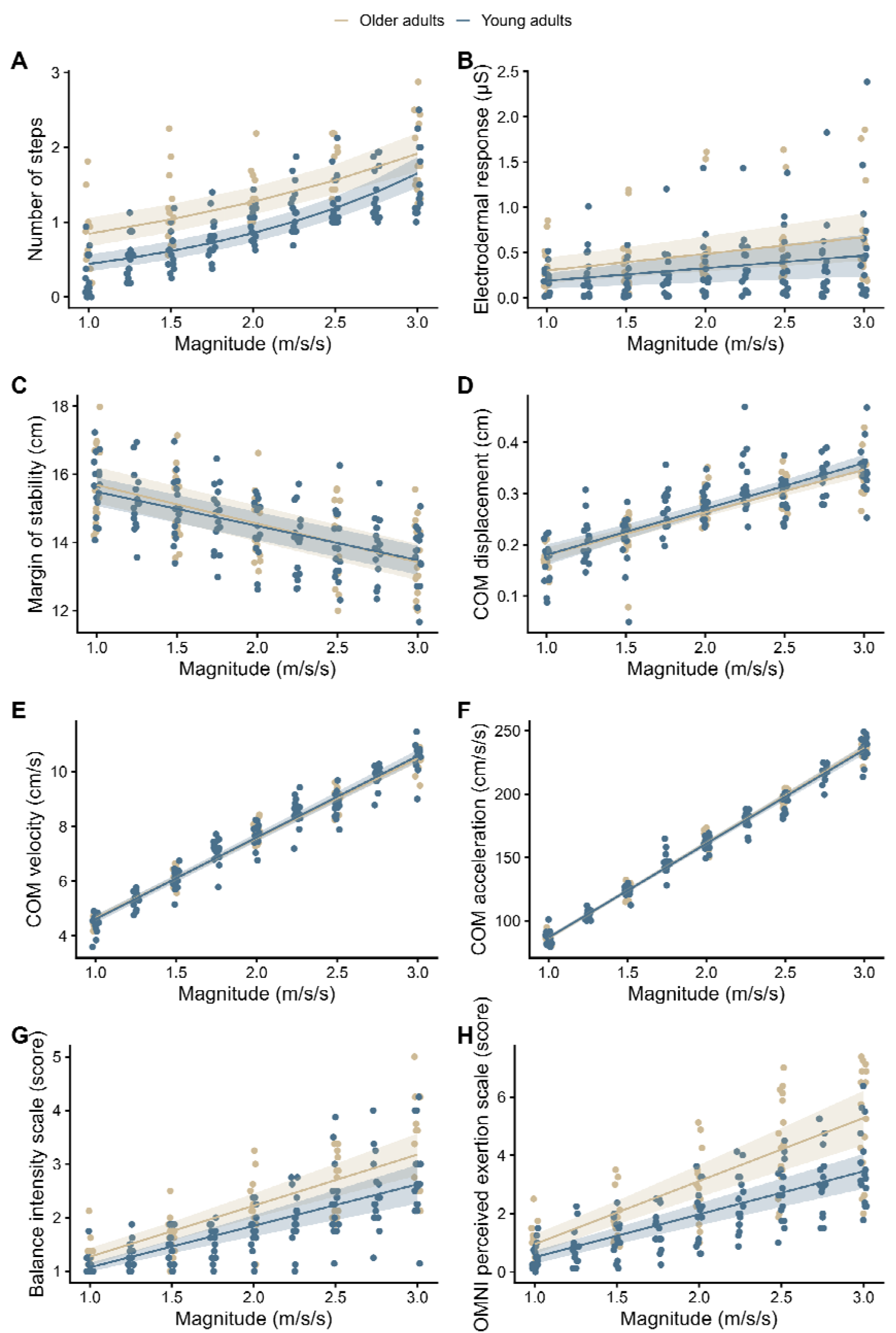
Relationships between perturbation magnitudes and candidate measures. Points show mean values for individual participants at each perturbation magnitude. Trend lines and confidence intervals (shaded area) are shown for each group.

Results of the comparison between groups and directions are shown in Figure 2.

**Figure 2:**
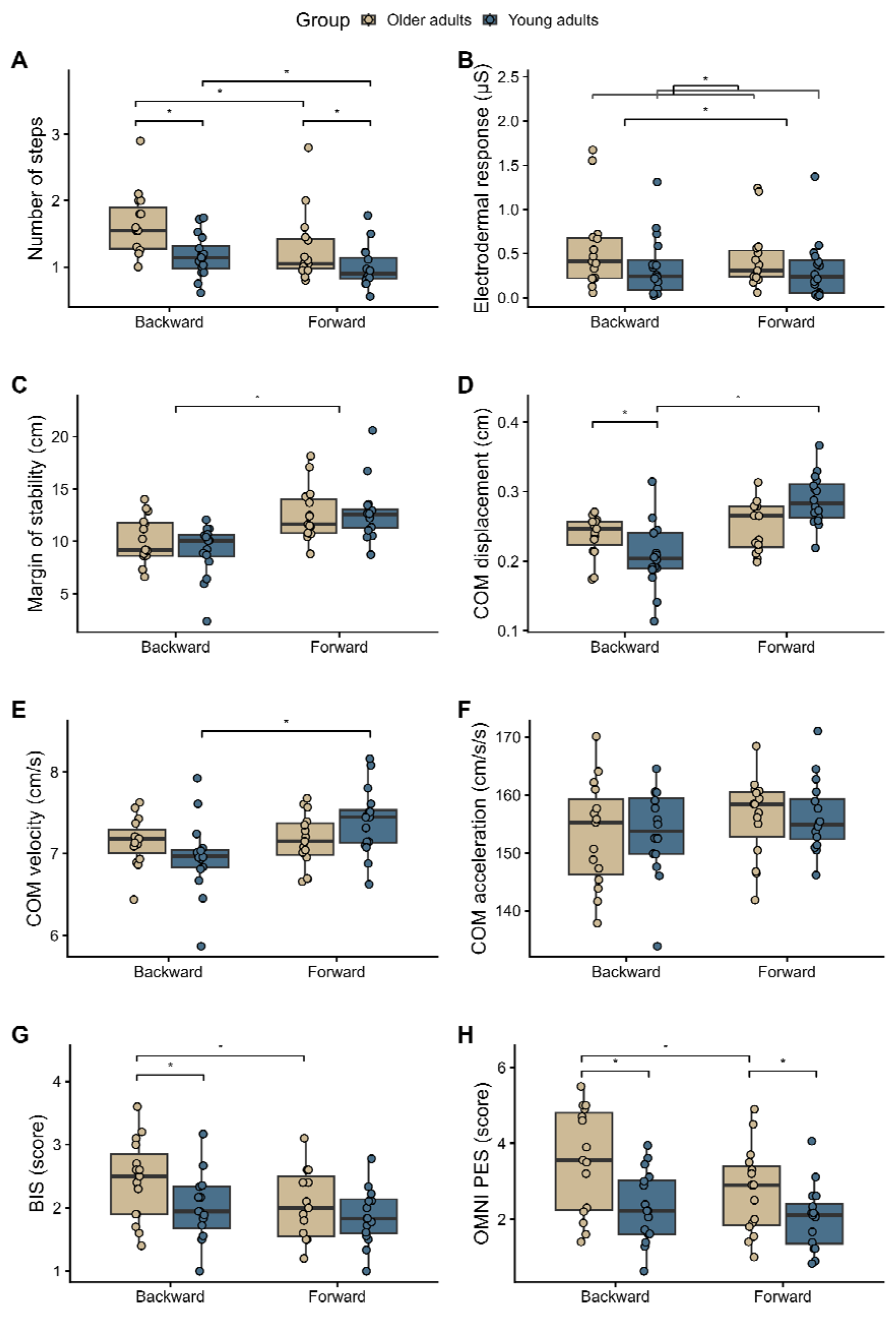
Comparison of measures between groups and directions. Points are individual data points for each participant. Significant differences are indicated with square brackets and asterisks. COM = centre of mass; BIS = Balance Intensity Scale; OMNI PES = OMNI perceived exertion scale score.

There was a significant group-by-direction interaction effect for number of steps (F_1,29_=4.68, p=0.039). Older adults took more steps than young adults when responding to forward (mean difference=0.31 [0.15, 0.48] steps] and backward falls (mean difference=0.51 [0.35, 0.67]). Furthermore, both groups took more steps when responding to backward than forward falls (older adult mean difference=0.37 [0.20, 0.55], younger adult mean difference=0.18 [0.007, 0.35]). There was no significant group-by-direction interaction effect (F_1,29_=3.55, p=0.070) for EDR. EDRs were significantly higher for older adults compared to young adults (mean difference=0.16 [0.11, 0.12] μS; F_1,29_=43.49, p<0.0001), and higher for backward-fall compared to forward-fall perturbations (mean difference=0.08 [0.02, 0.13] μS; F_1,29_=9.12, p=0.0052). There was no significant group-by-direction interaction effect (F_1,29_=0.54, p=0.47) or main effect of group (F_1,29_=0.16, p=0.69) for MOS. There was a significant main effect of direction for MOS (F_1,29_=15.03, p=0.0006); MOS was lower for backward-versus forward-fall perturbations (mean difference=3.1 [1.5, 4.8] cm). There were significant group-by-direction interaction effects for peak COM displacement (F_1,29_=9.27, p=0.0049) and velocity (F_1,29_=5.82, p=0.022). Post-hoc testing revealed that peak COM displacement and velocity were higher for young adults for forward falls compared to backward falls (displacement mean difference=0.09 [0.04, 0.13] cm; velocity mean difference=0.48 [0.13, 0.84] cm/s); there were no significant differences between directions for older adults. COM displacement was also higher for older adults than young adults for backward falls (mean difference =0.05 [0.002, 0.09] cm). There was no significant group-by-direction effect (F_1,29_=0.12, p=0.73), or main effects of group (F_1,29_=0.04, p=0.83) or direction (F_1,29_=1.47, p=0.24) for COM acceleration. There were significant group-by-direction interaction effects for BIS (F_1,29_=5.43, p=0.027) or OMNI PES scores (F_1,29_=5.17, p=0.031). BIS and OMNI PES scores were higher for backward falls than forward falls for older adults only (BIS mean difference=0.38 [0.15, 0.61]; OMNI mean difference=0.79 [0.37, 1.21]). BIS scores were higher for older adults than younger adults for backward falls only (mean difference=0.41 [0.21, 0.62]), whereas OMNI PES scores were higher for older than younger adults for both forward (mean difference=0.66 [0.25, 1.08] and backward falls (mean difference=1.15 [0.74, 1.56]).

## DISCUSSION

This study aimed to identify measures of absolute and relative RBT intensity. Using moving platform perturbations, where the absolute perturbation intensity is proportional to the peak platform acceleration, we found that all candidate measures were significantly related to peak platform acceleration for both groups of participants (young and older adults). However, only peak COM acceleration in the 100 ms after perturbation onset showed no evidence of significant differences between groups and directions, suggesting that this is a valid measure of absolute RBT intensity. The earliest responses to balance perturbations are typically initiated approximately 100 ms or later after perturbation onset (42). Therefore, COM motion during this early phase is related to features of the perturbation and is independent of the quality of the participants’ response (19).

Behavioural response (number of steps), EDR, and the OMNI PES significantly differed between groups and directions (for older adults only for the OMNI PES). This finding suggests that these measures reflect relative RBT intensity. Previously, in the absence of validated relative balance training intensity measures, some have applied the Rate of Perceived Exertion (43) to subjectively assess intensity of balance training (1,4,44). However, this scale is only valid for cardiorespiratory exercise (4,43). Lack of valid measures of balance training intensity have led others to more recently develop balance-specific scales to rate perceived challenge (25,45). In the current study, we demonstrated that one of these scales – the BIS – is valid for RBT. The Rate of Perceived Stability (RPS) scale has also been validated for anticipatory balance training (46,47). However, RPS may not be suitable for RBT as participants should score 10/10 on this scale (“about to fall: extremely challenged; have to step and/or grab support to keep balance”) during RBT. We also validated the OMNI PES for RBT, which has been previously validated for anticipatory balance training (28). The OMNI PES may be advantageous over the BIS as it is an 11-point, rather than a 5-point, scale, providing greater resolution for participant ratings. We also found that OMNI PES scores differed between directions for young adults. Therefore, the OMNI PES may be better able to differentiate differences in RBT intensity than the BIS.

Intensity rating scales may be difficult to use with people who have cognitive, communication, or language barriers. Even among people without cognitive, communication or language barriers, rating scales may be misused or misinterpreted (48). Furthermore, perceptions of intensity are subjective, and may be influenced by factors other than exercise intensity, such as age, sex and familiarity with the task (48,49). Therefore, it is valuable to complement participants’ perception of intensity with objective measures. Previous studies used frequency of single- or multi-step reactions (15,17,18), or a proportion of the multi-step threshold (16) to define RBT intensity; our study supports this approach for prescribing intensity of RBT. While the perturbation magnitudes used in the current study were not large enough to cause falls, others have shown that falls (into a safety harness) are more prevalent at higher perturbation magnitudes and more prevalent for people with stroke compared to controls without stroke at the same perturbation magnitude (50). It is reasonable, therefore, to assume that falling as a behavioural response, or needing assistance from a ‘spotter’ to avoid a fall, indicates a higher intensity than multi-stepping. In agreement with previous research showing that EDRs increase with increasing challenge or reactive (23) and anticipatory (51) balance tasks, and higher after balance perturbations for people with greater balance impairment (24), we also found that EDRs reflect relative RBT intensity.

Studies have varied FITT parameters to optimize exercise prescription for other types of exercise (52,53,e.g., for cardiorespiratory and resistance training;,54), revealing interdependence of FITT parameters; e.g., as intensity increases, frequency or time can decrease to achieve similar benefits (52,53). This research has contributed to exercise guidelines describing the optimal FITT for improving these components of fitness (2). Similar research is lacking for balance training, so optimal FITT cannot reported in exercise guidelines for balance training (1,2,55). Because FITT parameters are interdependent, intensity of balance training needs to be defined and measureable to facilitate research on optimal FITT. Clearly defined exercise intensity also facilitates determining a maximal intensity for some groups (e.g., novice exercisers or those with medical precautions), above which there may be an increased risk of injury or other adverse events (2,56). The current study findings may help to facilitate this future research on optimal FITT for, and safety of, RBT.

This study supports COM acceleration early after the perturbation as a measure of absolute RBT intensity, and number of steps, EDR, BIS, and OMNI PES as measures of relative RBT intensity. Future RBT studies should report how intensity of training was defined, and consider incorporating measures of behavioural response and participant perceptions of intensity (using either the BIS or OMNI PES) into training protocols. Future research should also explore objective measures of anticipatory balance training to facilitate developing guidelines for optimal balance training prescription.

## Acknowledgements

We acknowledge Nourali Ibrahim El Husseini Cardenas, Piufat Wong, Isabel Yeh, Laura McPhail, and Chiara Tumminieri for their assistance with data collection.

